# Cortical tracking of continuous speech under bimodal divided attention

**DOI:** 10.1101/2022.10.29.514344

**Authors:** Zilong Xie, Christian Brodbeck, Bharath Chandrasekaran

## Abstract

Speech processing often occurs amidst competing inputs from other modalities, e.g., listening to the radio while driving. We examined the extent to which *dividing* attention between auditory and visual modalities (bimodal divided attention) impacts neural processing of natural continuous speech from acoustic to linguistic levels of representation. We recorded electroencephalographic (EEG) responses when human participants performed a challenging primary visual task, imposing low or high cognitive load while listening to audiobook stories as a secondary task. The two dual-task conditions were contrasted with an auditory single-task condition in which participants attended to stories while ignoring visual stimuli. Behaviorally, the high load dual-task condition was associated with lower speech comprehension accuracy relative to the other two conditions. We fitted multivariate temporal response function encoding models to predict EEG responses from acoustic and linguistic speech features at different representation levels, including auditory spectrograms and information-theoretic models of sublexical-, word-form-, and sentence-level representations. Neural tracking of most acoustic and linguistic features remained unchanged with increasing dual-task load, despite unambiguous behavioral and neural evidence of the high load dual-task condition being more demanding. Compared to the auditory single-task condition, dual-task conditions selectively reduced neural tracking of only some acoustic and linguistic features, mainly at latencies >200 ms, while earlier latencies were surprisingly unaffected. These findings indicate that behavioral effects of bimodal divided attention on continuous speech processing occur not due to impaired early sensory representations but likely at later cognitive processing stages. Crossmodal attention-related mechanisms may not be uniform across different speech processing levels.

## Introduction

Speech processing often occurs amidst competing inputs from other sensory modalities, e.g., listening to the radio while driving. In such situations, listeners must allocate attention across modalities to effectively select the most relevant information within a modality. This raises the question of whether and how *dividing* attention between modalities (e.g., audition and vision; bimodal divided attention) affects the processing of natural continuous speech.

Resource-based theoretical frameworks have been invoked to scaffold the understanding of mechanisms governing crossmodal attention (Wahn & König, 2017). Two contrastive resource-based accounts (modality-specific versus supramodal) yield different hypotheses regarding the effects of bimodal divided attention on continuous speech processing. Per the *modality-specific* account, each sensory modality is allocated a limited pool of attentional resources, and these pools of attentional resources operate independently of each other (Alais et al., 2006; Arrighi et al., 2011; Duncan et al., 1997; Keitel et al., 2013; Parks et al., 2011; Porcu et al., 2014). In contrast, per the *supramodal* account, different sensory modalities share a central, limited pool of attentional resources. The availability of resources to one modality is inversely related to the amount of resources used by other modalities (Broadbent, 1958; Ciaramitaro et al., 2017; Klemen et al., 2009; Macdonald & Lavie, 2011; Molloy et al., 2015).

Empirical evidence regarding bimodal divided attention effects on speech processing primarily comes from experimenter-constrained tasks (e.g., Gennari et al., 2018; Kasper et al., 2014; Mattys et al., 2009, 2014; Mattys & Palmer, 2015; Mattys & Wiget, 2011). Many studies have shown the detrimental effects of bimodal divided attention on the acoustic processing of simplified, controlled speech stimuli (e.g., syllable or single words) (Gennari et al., 2018; Mattys et al., 2014; Mattys & Palmer, 2015; Mattys & Wiget, 2011), which is consistent with the *supramodal* account of attention. Speech processing entails mapping acoustic features into linguistic representations of increasing complexity (Brodbeck & Simon, 2020; Hickok & Poeppel, 2007), raising the question of how bimodal divided attention affects linguistic representations beyond acoustic processing. Behavioral studies with simple speech stimuli indicate that reduced acoustic processing under bimodal divided attention may lead to compensatory changes manifested by increased reliance on higher-order linguistic knowledge during auditory lexical perception (Mattys et al., 2009). However, to date, there is a lack of a systematic and holistic analysis of divided attention-related changes across different levels (acoustic-to-linguistic) of natural continuous speech processing, which is distinctly different from processing simple speech stimuli (Gaston et al., 2022; Hamilton & Huth, 2020).

Here, we assessed electroencephalography (EEG) to provide a systematic and holistic analysis of the acoustic and linguistic processing of continuous speech (Brodbeck & Simon, 2020; Gillis et al., 2022). The continuous speech paradigm uses the multivariate temporal response function approach (Crosse et al., 2016; Ding & Simon, 2012) to predict neural responses from a combination of hypothesis-driven acoustic and linguistic properties of continuous speech. The predictive power of each speech property is used to quantify the corresponding processing levels (Brodbeck & Simon, 2020; Gillis et al., 2022). The spectro-temporal acoustic properties included envelope-based spectrogram and acoustic onset spectrogram. The linguistic properties included measures of informativeness (surprisal and entropy) based on the information-theoretic framework (Brodbeck et al., 2018). Prior work suggests that both acoustic and linguistic representations are strongly modulated by *selective* attention, within the auditory modality and across modalities. Attentional effects are disproportionality more robust on the linguistic representations than acoustic-based representations (Brodbeck et al., 2018, 2020).

Here we integrated the continuous speech paradigm with an audiovisual dual-task paradigm to examine the effects of bimodal divided attention on the acoustic and linguistic processing of continuous speech. In the dual-task paradigm, participants performed a challenging primary visuospatial task that imposed low or high cognitive load while listening to audiobook stories as a secondary task. The two dual-task conditions were contrasted with an auditory single-task condition in which participants attended to the story while ignoring visual stimuli. We hypothesized that compared to the auditory single-task condition, dual-task conditions would lead to reduced acoustic and linguistic representations of continuous speech, especially at high cognitive load. However, we hypothesized that linguistic representations may be affected to a relatively greater extent based on evidence from the literature on selective attention. These hypotheses are aligned with the *supramodal* account of crossmodal attention.

## Materials and Methods

### Experimental design

Bimodal divided attention was manipulated via a dual-task paradigm. Specifically, participants performed a primary visuospatial *n*-back task of varying (high or low) cognitive load (Jaeggi et al., 2007) while listening to continuous speech as a secondary task. We designated the visual task as the primary task to maximize the chance of observing the bimodal divided attention effects on continuous speech processing. The cognitive load of the dual-task paradigm was manipulated via 3- and 0-back tasks on the visuospatial stimuli (blue squares; Figure 1A and 1B). The dual-task conditions were contrasted with an auditory single-task condition (Figure 1C), in which participants explicitly attended to the auditory stimuli while ignoring the visual stimuli. To obtain a behavioral measure for the auditory task, participants were instructed to respond to two multiple-choice comprehension questions on the story segments at the end of each trial. Detailed task instructions are presented in the section on *Experimental procedure*.

Each task condition consisted of 15 trials of visual stimuli paired with 15 unique story segments and were presented in separate blocks. The order of the story segments was fixed and identical across participants in order to maintain the continuity of the storyline. The order of task conditions was counterbalanced across participants. Each trial of visual stimuli ended later than the corresponding story segment. Such offset gaps were not significantly different across task conditions [*F*(2, 42) = .01, *p* = .99]. The experiment was controlled with E-Prime 2.0.10 (Schneider et al., 2002).

**Figure 1.**
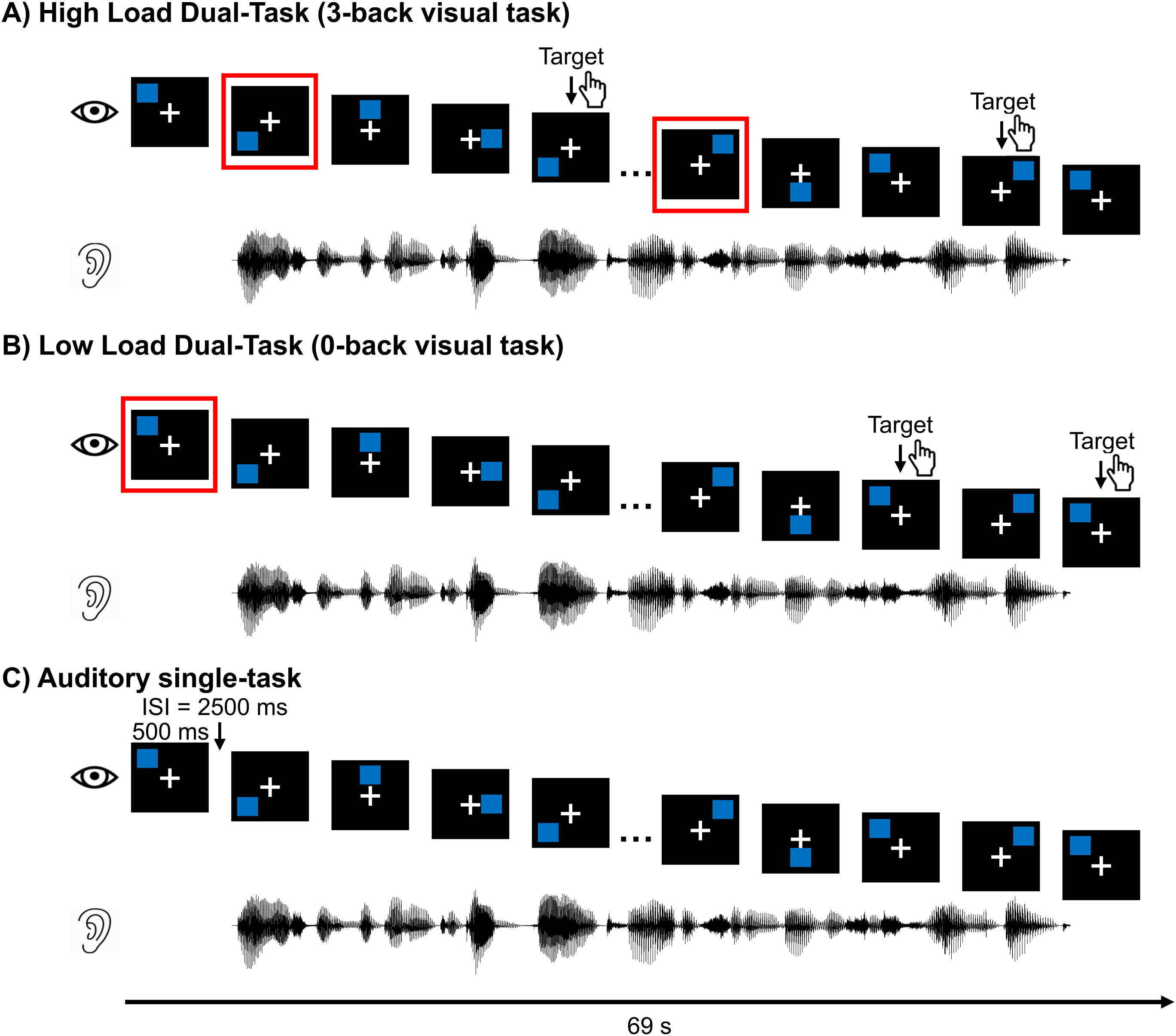
Trial design illustrations for (A) high load dual-task (3-back visual ask), (B) low load dual-task (0-back visual task), and (C) auditory single-task condition. In the two dual-task conditions, the primary task was to respond to the visual stimuli and the secondary task was to attend to auditory stimuli (story segments of about 60 seconds). In the high load condition (A), participants responded only when the current blue square matched the one 3 positions back (examples highlighted in red squares). In the low load condition (B), participants responded only when the current blue square matched the first square in each trial (highlighted in the red square). In the auditory single-task condition (C), participants were instructed to attend to the auditory stimulus and ignore the visual stimuli. At the end of each trial, participants responded to two multiple-choice comprehension questions for the story segments. ISI: interstimulus interval.

### Participants

Adult native American English speakers (N = 18) were recruited from the Austin, Texas, community. Data from one participant were excluded due to technical problems. Data from another participant were excluded because their story comprehension accuracy was lower for the auditory single-task condition (66.67%) than the two dual-task conditions (73.37% for low load and 76.67% for high load). We interpreted this result as that this participant did not understand or follow the task instructions. The final sample consisted of sixteen participants (18 to 23 years old; 11 females, five males; 14 right-handed and two left-handed). The sample size was selected based on prior work examining the effects of bimodal attention on the neural processing of speech stimuli (e.g., Gennari et al., 2018; Kasper et al., 2014). Previous studies have shown that music training can influence speech processing (e.g., Bidelman & Alain, 2015). Therefore, we recruited only participants without a history of or significant formal music training (<= four years of continuous training, not currently practicing). All participants had normal air and bone-conduction audiometric thresholds, defined as <= 20 dB hearing level for octave frequencies from 0.25 to 8 kHz. The thresholds were measured via an Interacoustics Equinox 2.0 PC-Based Audiometer. Additional inclusion criteria are as follows: no history of psychological or neurological disorders, no use of neuropsychiatric medication, and having normal or corrected-to-normal vision. Before the experiment, all participants provided written, informed consent. Participants received monetary compensation for their participation. The Institutional Review Board at the University of Texas at Austin approved the experimental protocols.

### Stimuli and apparatus

The stimuli were composed of visual and auditory materials. The visual stimuli (Figure 1) were blue squares at one of eight loci around a white fixation cross in the center of a black screen, adapted from Jaeggi et al. (2007). The duration for individual squares was 500 ms, and the interval between consecutive squares was 2500 ms. Twenty-three squares were included in a trial, lasting 69 seconds. The stimuli were displayed on a VIEWPixx/EEG LCD monitor with a scanning LED-backlight design [29.1 cm (height) × 52.2 cm (width); display resolution: 1920 × 1080; refresh rate: 120 Hz] at an ~110 cm distance from participants’ eyes.

The auditory stimuli were English audiobook stories selected from a classic work of fiction, *Alice’s Adventures in Wonderland* (Chapters 1-7, http://librivox.org/alices-adventures-in-wonderland-by-lewis-carroll-5). The audiobook was narrated by an adult male American English speaker at a sampling rate of 22.05 kHz. The chapters were divided into 45 segments (each ~60 seconds long). Each segment began where the story ended in the previous segment. In each segment, silent periods of more than 500 ms were shortened to 500 ms. The story stimuli were presented diotically via insert earphones (ER-3; Etymotic Research, Elk Grove Village, IL) to the participants at a 70 dB sound pressure level. A trial of visual stimuli (23 blue squares) was presented concurrently with each story segment, with the segment beginning later (3 seconds after the onset of the visual trial) and ending earlier relative to the visual trial.

### Experimental procedure

#### High and low load dual-task

The cognitive load of the dual-task conditions was manipulated via the visual task. For the high load condition, the visual task required participants to respond when the current blue square matched the one three-position back in the sequence (i.e., 3-back task, Figure 1A). For the low load condition, the visual task required participants to respond when the current blue square matched the first square in the sequence (i.e., 0-back task, Figure 1B). We randomized the location of the first square across trials. Matched squares were treated as targets, and unmatched ones were non-targets. Note that targets could appear only starting from the fourth square in the sequence for a given trial in the 3-back task. In other words, targets would be among the last 20 squares in the sequence on a given trial. We designed the 0-back task to match that. Six of the last 20 squares were set as targets for both task conditions, and the remaining 14 were non-targets. The target locations were randomized across trials.

Participants responded to the targets by pressing buttons on a game controller. Participants were told that speed and accuracy were equally important. Participants were required to rest their fixations on a white cross in the middle of the screen. To encourage engagement, accuracy feedback on the visual task was displayed after their responses. The number of button presses was not significantly different between 3- and 0-back visual tasks [*t*(15) = .96, *p* = .36]. After the visual task, participants responded to two multiple-choice comprehension questions for the auditory stories. Participants had unlimited time to respond to the story questions. No feedback was provided after their responses.

Critically, to manipulate the priority of the auditory and visual tasks, participants were instructed to focus primarily on the visual task and attend to the auditory stimulus as a secondary task. They were explicitly told that their data could not be used if their performance on the visual task was poor.

#### Auditory single-task

In this condition, participants were instructed to focus on the story segments and ignore the visual stimuli (Figure 1C). Participants were required to keep their eyes open and rest their fixations on a white cross in the middle of the screen. At the end of each trial, participants responded to two multiple-choice questions to assess their comprehension of the story segments. Participants had unlimited time to respond to questions. Visual feedback about the accuracy of the story question was displayed following their responses.

### Electrophysiological data acquisition and preprocessing

#### Acquisition

The experiment was conducted in a dark, acoustically shielded booth. Participants were seated in a comfortable chair during tasks. Electroencephalography (EEG) data were recorded using the Easycap recording cap (www.easycap.de) with 64 actiCAP active electrodes (Brain Products, Gilching, Munich, Germany) at a sampling rate of 5 kHz. The electrode locations were determined according to the extended 10-20 system (Oostenveld & Praamstra, 2001), with a common ground at the Fpz electrode site and TP9 as the reference. The electrode impedances were below 20 kΩ.

The EEG data were acquired using BrainVision actiCHAmp amplifier (Brain Products, Gilching, Munich, Germany) linked to BrainVision Pycorder software 1.0.7.

#### Preprocessing

The EEG data were preprocessed offline in MNE-Python (Gramfort et al., 2013), and the deconvolution analysis was implemented with the Eelbrain package (Brodbeck et al., 2021). The data were re-referenced to the average of the electrodes TP9 and TP10, and then band-pass filtered from 1 to 15 Hz using a zero-phase overlap-add finite impulse response filter (hamming window) with default settings in MNE-Python. Independent component analysis was applied to EEG data combined across the three task conditions in individual participants using the extended-infomax algorithm. Artifact-related components (mainly ocular artifacts) were identified according to the topographical distribution and time course via visual inspection and then removed. After that, the EEG data were segmented into epochs that were time-locked to the onsets of corresponding story segments, and then downsampled to 100 Hz. The maximum possible duration of the epochs was set to 61 seconds.

### Assessing neural tracking of visual and auditory stimuli

To assess the neural representation of speech, we used the multivariate temporal response function (mTRF) approach (Crosse et al., 2016; Ding & Simon, 2012). In this approach, the EEG signal is predicted using time-delayed multiple regression. We first generated several visual, acoustic, and linguistic models (see below). Each model was used to define several predictor variables, each implementing a specific linking hypothesis for predicting brain activity from the corresponding model. We then tested the predictive power of different combinations of predictor variables to evaluate which acoustic and linguistic models are associated with predictive power for the EEG data. Each predictor variable thus operationalizes a hypothesis that EEG responses are modulated by a given property of the speech signal, which would indicate neural representations arising from a corresponding acoustic or linguistic model. Figure 2 displays an example of each predictor variable. In the following paragraphs, we provide more detailed descriptions.

**Figure 2.**
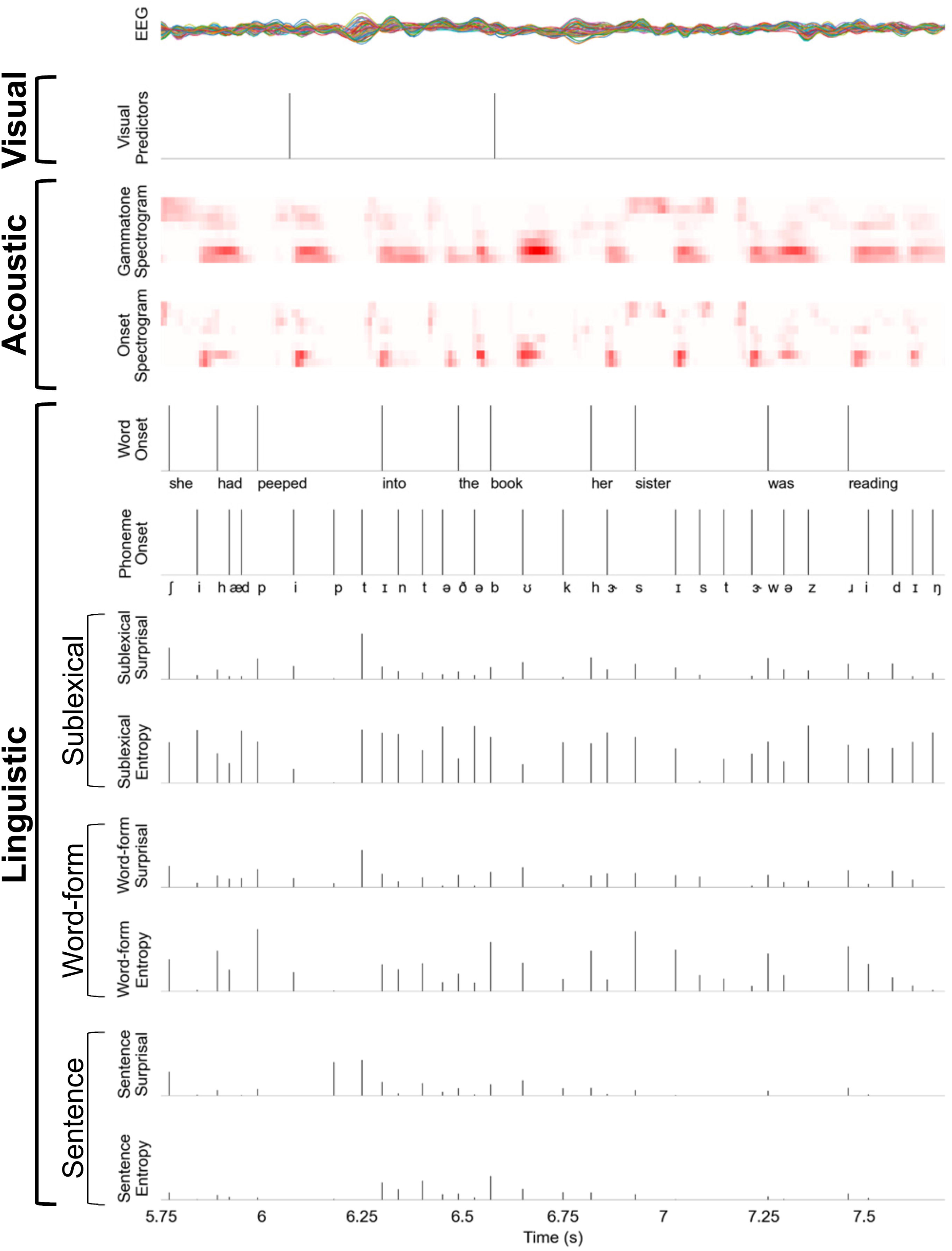
An excerpt of raw EEG responses from all 64 electrodes (top row) and the predictor variables (subsequent rows) used to model the EEG responses. Note that visual predictors consist of a separate one-dimensional array with impulses for onsets and offsets of the blue squares. They are combined into a single predictor in this example for illustration purposes.

#### Visual model

Because the visual stimuli were temporally sparse, visual responses were modeled analogously to a visual evoked potential. The visual predictor was a one-dimensional time series with an impulse at the onsets and offsets of the blue squares. We did not separate predictors for targets and non-targets because this study was not intended to explore differences in neural processing of visual targets and non-targets, and thus there were not enough targets to estimate stable responses.

#### Acoustic model

The acoustic model was designed to assess EEG responses related to representations of acoustic spectro-temporal features. All acoustic predictors were derived from 256-band gammatone-based spectrograms of the speech stimuli, with cut-off frequencies from 0.02 to 5 kHz. The 256-band spectrograms were downsampled to 1 kHz and scaled with an exponent of 0.6. A *spectrogram* predictor was then created by summing the 256-band spectrograms in eight logarithmically spaced frequency bands. In addition, an *onset spectrogram* predictor was defined to detect and control for representations of acoustic edges. These were generated using an auditory edge detection model (Brodbeck et al., 2020; Fishbach et al., 2001) and applied to each frequency band of the 256-band spectrograms. The onsets across these 256 bands were also summed into eight logarithmically spaced frequency bands to generate 8-band onset spectrogram predictors.

#### Linguistic models

Linguistic processing was assessed using information-theoretic models. These models assume that listeners maintain predictive models of speech, which can be linked to brain activity through surprisal and entropy measures (Brodbeck et al., 2018). Previous work suggests that listeners maintain multiple such predictive models, differing in complexity, in parallel (Brodbeck et al., 2022). The predictive models were all defined over phoneme sequences, determined for each stimulus via forced alignment using the Montreal Forced Aligner (MFA) (McAuliffe et al., 2017). The predictors based on the specific information-theoretic models (described in subsequent sections) all consisted of time series with an impulse of variable size at each phoneme onset. In order to provide a control for responses related to linguistic segmentation, two additional predictors were included: A *word onset* predictor with a unit (value of 1) impulse at the onset of each word-initial phoneme and a *phoneme onset* predictor with a unit impulse at the onsets of all other phonemes.

##### Sublexical model

The sublexical model assumes that listeners predict upcoming phonemes or speech sounds based on a local context, consisting of only a few preceding sounds. To implement such a model, all sentences from the SUBTLEX-US corpus (Keuleers et al., 2010) were transcribed into phoneme sequences without word boundary markers, and a 5-gram model (Heafield, 2011) was trained on these phoneme sequences. This model was then applied to the experimental stimuli to compute a probability distribution over phonemes at each phoneme position, conditional on the four preceding phonemes. This distribution was used to calculate a *sublexical surprisal* predictor (the surprisal of encountering phoneme *ph_k_* at position *k* in the stimulus is −*log*_2_ (*p*(*ph_k_*|*context*)), and a *sublexical entropy* predictor (the entropy at phoneme position *ph_k_* reflects the uncertainty about what the next phoneme, *ph_k_*_+1_, will be 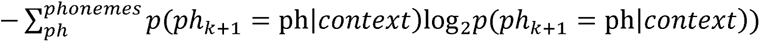. Surprisal is a measure of the amount of new information provided by a stimulus; a response to sublexical surprisal is thus evidence that listeners integrate information on a sublexical level. A response to entropy additionally suggests that listeners create a probabilistic expectation about future input before encountering the phoneme (Pickering & Gambi, 2018). A response to either of those predictors would provide evidence that listeners maintain a sublexical language model.

##### Word-form model

The word-form model aims to predict the surface form of the word that is currently being heard, but without access to any information preceding the word. To implement this model, a phonological lexicon was generated by combining pronunciations from the MFA English dictionary and the Carnegie Mellon University Pronouncing Dictionary (http://www.speech.cs.cmu.edu/cgi-bin/cmudict). The word-form model was implemented based on the cohort model of word recognition (Brodbeck et al., 2018; Marslen-Wilson, 1987). Each word was assigned a prior probability based on its count frequency in the SUBTLEX corpus (Keuleers et al., 2010), substituting a count of 1 for missing words. For each word in the speech stimuli, the cohort model was then implemented by starting with the complete lexicon and, for each subsequent phoneme of the word, incrementally removing words that were not compatible with that phoneme in that position. The changing probability distribution over the lexicon was then used to define two predictors, each with a value at each phoneme position: *phoneme surprisal* (log inverse of the posterior probability of the phoneme given the preceding phonemes) and *cohort entropy* (Shannon entropy over all words currently in the cohort, 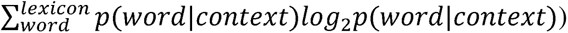. This model implements the hypothesis that listeners recognize words using a probabilistic model that takes into account all the information since the last word boundary (i.e., where the word started), but that does not further take into account any context when considering possible word forms as candidates.

##### Sentence model

The sentence model is very similar to the word-form model, with the only difference being that the prior expectation for each word is modulated by the sentence context. To implement this, a lexical 5-gram model (Heafield, 2011) was trained on the whole SUBTLEX-US database (Keuleers et al., 2010). This 5-gram model was used to set the prior probability for each word in the lexicon based on the preceding four words before applying the procedure described for the word-form model above. The same two linking hypotheses were used to define predictor variables (*phoneme surprisal* and *cohort entropy*). The sentence model implements the hypothesis that listeners use a wider context including multiple words, when modulating their phoneme-by-phoneme perception and expectations.

#### Estimation of neural tracking

We used forward encoding mTRF models to predict EEG responses from the predictors described above. The mTRF models were fitted to the EEG responses at individual electrodes using the boosting algorithm implemented in the Eelbrain toolbox (Brodbeck et al., 2021). The predictive power of the mTRF models was evaluated by how accurately they could predict EEG responses from novel trials of the same condition. This was quantified through the *z*-transformed Pearson’s correlation coefficient between predicted and actual EEG responses (i.e., prediction accuracy). Higher prediction accuracy indicates better neural tracking of the corresponding predictor.

The mTRFs were estimated separately for each subject and condition using a 5-fold cross-validation strategy. First, the trials were divided into 5 test sets. For each test set, EEG responses were predicted from the average of 4 mTRF models, estimated from the remaining 4 datasets, each with 3 of the remaining 4 sets serving as training data, and one as validation set. The mTRFs were generated from a basis of 50 ms Hamming windows with stimulus-EEG lag from −100 to 500 ms (window center). The mTRFs were estimated jointly for all predictors with coordinate descent to minimize the ℓ_1_ error. After each step, the change in error was evaluated in the validation set, and if there was an increase in error, the TRF for the predictor responsible for the change was frozen (in its state before the change). This continued until the whole mTRF was frozen. A single measure of prediction accuracy (fisher *z*-transformed correlation between predicted and measured response, see above) was calculated after concatenating the predicted responses from the 5 test sets. For analysis of the response functions, the mTRFs were averaged across all the test sets. For the visual predictor, the TRFs to onsets and offsets were combined for an effective response function with lags from −100 to 1000 ms relative to visual stimulus onset (because the visual stimulus always lasted exactly 500 ms).

To estimate the neural tracking of a given predictor (or a combination of predictors), we calculated the change in prediction accuracy (i.e., Δ*z*) when the predictor(s) of interest was(ere) removed from the full model that included all the visual, acoustic, and linguistic predictors. This procedure tests for variability in the responses that can be *uniquely* attributed to the predictor(s) under investigation and cannot be explained by any other predictors. Such a strong test is warranted because different properties of natural, connected speech tend to be correlated in time. Note that the analysis of the mTRFs themselves could not implement such strict control, and thus we cannot exclude the possibility that response functions include components that are confounded with other, correlated speech features. For this reason we focus our interpretation on tests of predictive power more than mTRF comparisons.

### Statistical analysis

All statistical analyses, if unspecified, were implemented in the R software (version 4.2.1; Team, 2022).

First, we examined the effect of task condition (auditory single-task, or low or high load dual-task) on behavioral performance, and neural visual, acoustic, and linguistic processing separately. A paired T-test (two-sided), or one- or two-way repeated-measures analysis of variance (ANOVA), whichever was appropriate, was performed with an alpha level of .05. The reported *p* values of those analyses were adjusted using the False Discover Rate (FDR) method (Benjamini & Hochberg, 1995). We also calculated effect sizes [Cohen’s *d* for T-tests and partial eta squared (η^2^_p_) for ANOVAs] and Bayes Factors (BF). The Bayes Factors were computed using appropriate functions from the R package ‘BayesFactor’ (version 0.9.12.4.4; Morey et al., 2022). Post hoc analysis, if necessary, was performed using paired T-tests (two-sided). FDR-corrected *p* values, effect sizes (Cohen’s *d*), and Bayes Factors (BF) were reported. More analysis details are provided in the following paragraphs.

Behavioral performance was quantified by three measures, including the accuracy and reaction time (RT) for the visual task and the accuracy for the auditory task. Visual accuracy was calculated as the difference in hit rates (i.e., correctly responding to a target) and false alarm rates (i.e., identifying a non-target as being a target). Visual RT was calculated as the median RT for hits only (Jaeggi et al., 2007; Snodgrass & Corwin, 1988). Auditory accuracy was calculated as the percentage of correctly answered story questions.

The extent of neural visual processing was determined using a mass-univariate analysis, comparing the predictive power (*z*) between the full model and a model missing the visual predictor. For this, we averaged the prediction accuracy for visual predictors across task conditions at individual electrodes and tested whether the averaged difference in prediction accuracy (Δ*z*) was greater than zero using a mass-univariate, one-sample T-test (one-sided). This was implemented in the Eelbrain package. The mass-univariate test was a cluster-based permutation test, using a *t*-value equivalent to uncorrected *p* ≤ 0.05 as the cluster-forming threshold. A corrected *p*-value was computed for each cluster based on the cluster-mass statistic in a null distribution from 10,000 permutations (or a complete set of all possible permutations, in cases where this was fewer than 10,000) (Maris & Oostenveld, 2007). We reported the largest *t* value from the cluster, i.e., *t_max_*, as an estimate of effect size (Brodbeck et al., 2018). Neural acoustic and linguistic processing were analyzed in the same manner.

We followed each of these analyses by examining the extent to which task conditions modulated neural tracking of individual predictors, or subsets of predictors. To this end we used the significant cluster from the mass-univariate analysis as region of interest (ROI) to extract Δ*z* values averaged across the electrodes in the cluster, but for each condition separately. Regarding neural acoustic processing, we examined the spectrogram and onset spectrogram predictors separately. Regarding neural linguistic processing, we conducted three sets of analyses to examine individual linguistic predictors. First, a two-way repeated-measures ANOVA was performed to examine the effects of context levels (sublexical, word-form, and sentence) and task condition on prediction accuracy. Second, a two-way repeated-measures ANOVA was performed to examine the effects of predictor type (entropy and surprisal) and task condition on prediction accuracy. Third, a one-way repeated-measures ANOVA was performed to examine the effect of task condition on the prediction accuracy of word onsets. Further, if a significant effect of task condition was observed from any of those analyses, we conducted follow-up analyses to examine whether task conditions eliminated neural tracking of the corresponding predictor(s) by testing whether the prediction accuracy at individual task conditions was greater than zero using one-sample T-tests (one-sided).

Finally, we examined the effect of task conditions on the mTRFs from predictors which showed significant task conditions effects on prediction accuracy. The predictors include visual predictors, onset spectrogram, three context levels (sublexical, word-form, and sentence), and two predictor types (entropy and surprisal). We calculated the global field power (GFP) of mTRFs across the corresponding ROI from the above analyses of prediction accuracy. We then compared the GFP of mTRFs between task conditions using mass-univariate paired T-tests (two-sided). The mTRF analyses were implemented in the Eelbrain package with default parameters except for the analysis time window. For visual predictors, we concatenated the mTRFs for visual onsets and offsets to analyze the response to visual stimuli as a whole. For the onset spectrogram, we averaged the mTRFs across the eight frequency bands. The analysis time window was 0 to 1000 ms for visual predictors and 0 to 450 ms for auditory predictors.

## Results

Table 1 summarizes the key findings regarding the effect of task condition on behavioral performance, and neural visual, acoustic, and linguistic processing.

**Table 1.**
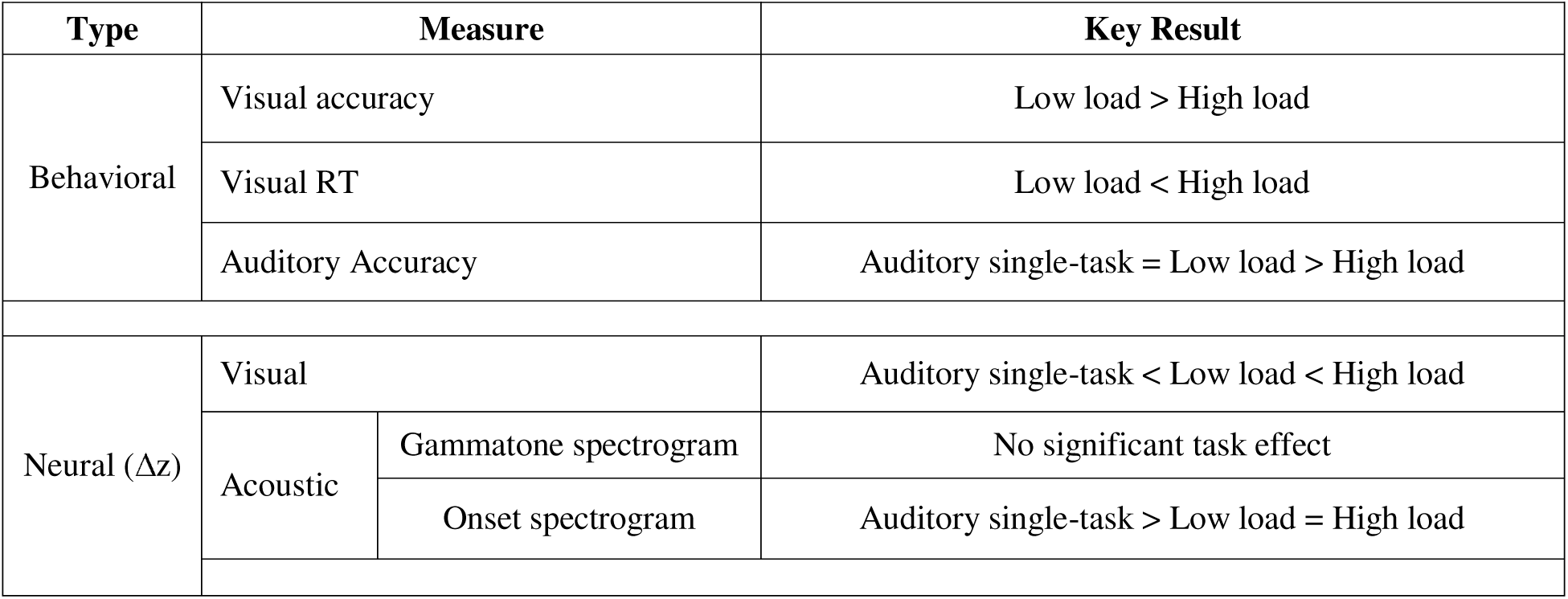
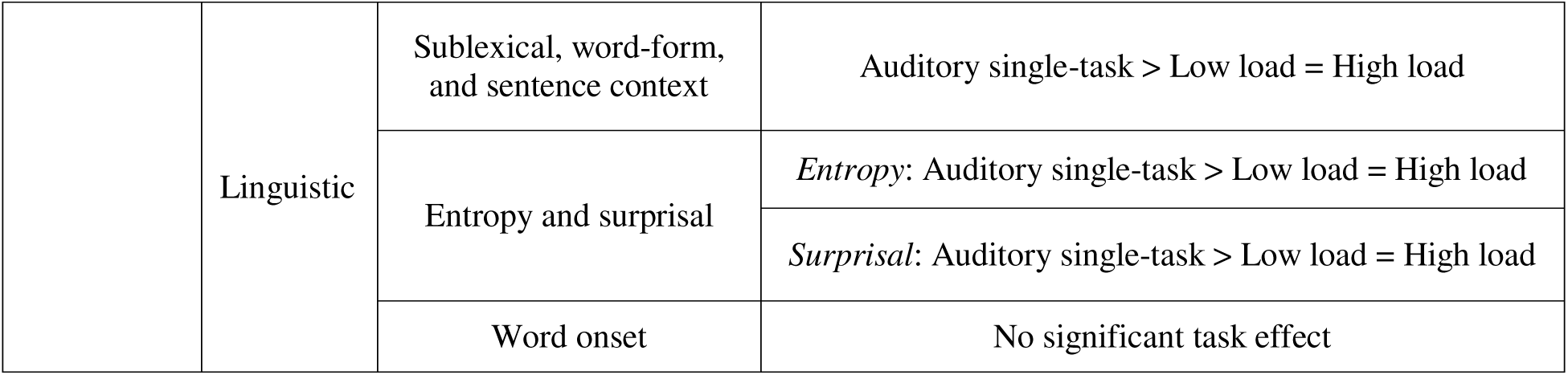
Task Effects on Continuous Speech Processing

### Divided attention and visual load impair behavioral performance

Figure 3A and 3B display the accuracy and RT of the visual task for individual participants. Compared to the low load (0-back) condition, the high load (3-back) condition was associated with lower accuracy [low load: mean = 99.54% (*SD* = 0.82) vs. high load: mean = 63.31% (*SD* = 21.85), *t* (15) = 6.60, *p* < .001, Cohen’s d = 1.65, BF = 2.59 × 10^3^] and slower RT [low load: mean = 453.11 ms (*SD* = 67.54) vs. high load: mean = 785.24 ms (*SD* = 233.41), *t* (15) = −5.33, *p* < .001, Cohen’s d = 1.33, BF = 330.30]. These results confirmed that the manipulation of cognitive load in the visual task was successful.

**Figure 3.**
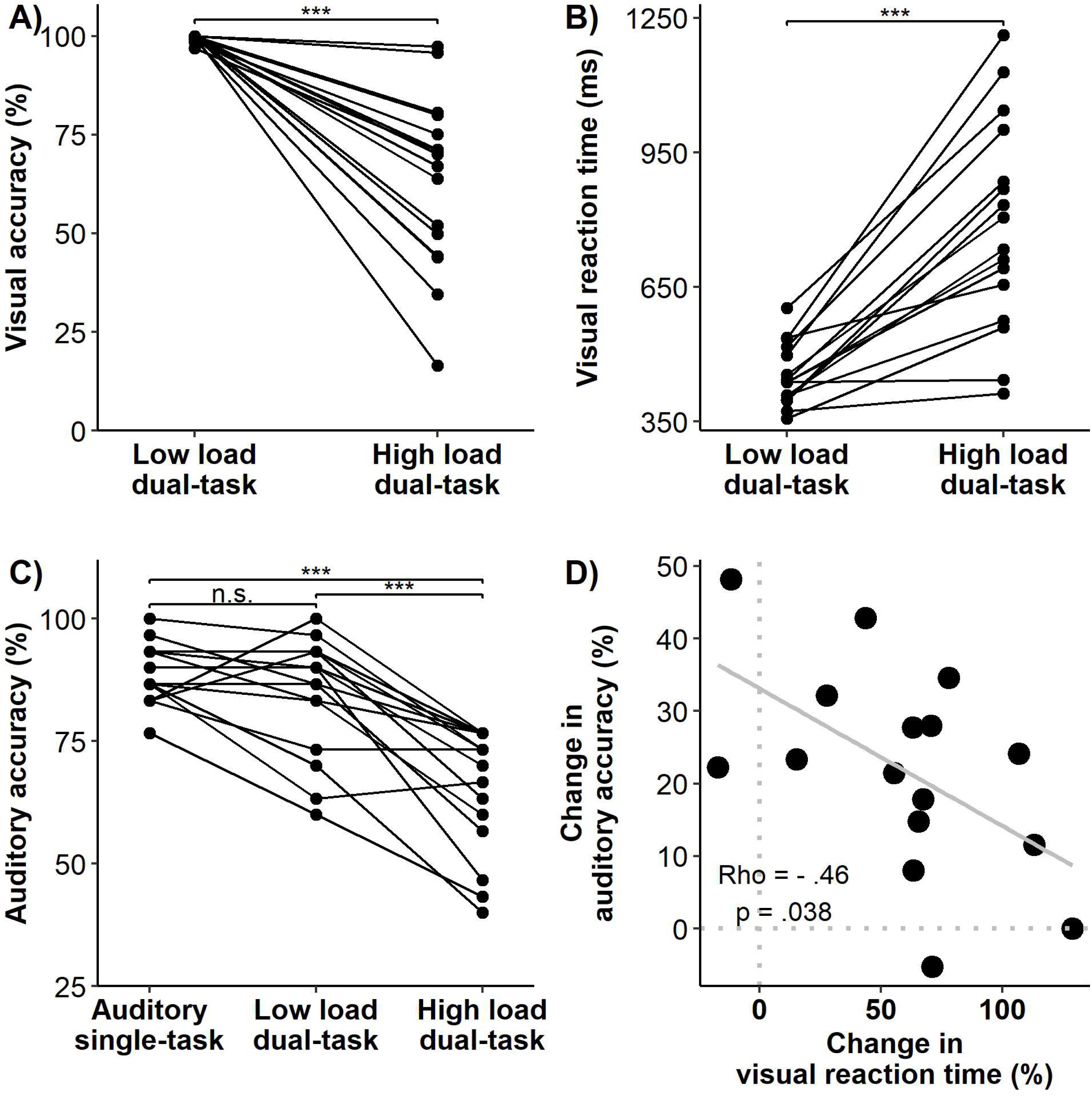
Behavioral performance on visual and auditory tasks. (A) Accuracy on the low load (0-back) and high load (3-back) visual tasks, which was calculated as the difference in hit rates (i.e., correctly responding to a target) and false alarm rates (i.e., identifying a non-target as being a target). (B) Reaction time (RT) on the low load (0-back) and high load (3-back) visual tasks, which was calculated for hits only. (C) Accuracy on the auditory task, which was calculated as the percentage of correctly answered story questions. Individual lines in (A) to (C) denote individual participants (n = 16). (D) Correlation between the change in auditory accuracy [i.e., (low load – high load)/low load] and the change in visual RT [i.e., (high load – low load)/low load]. The gray line is the linear regression line. N.s. *p* > .05, *** *p* < .001.

Figure 3C displays the auditory task accuracy for individual participants. The mean accuracy was 88.96% (*SD* = 5.93) in the auditory single-task condition, 84.58% (*SD* = 11.86) in the low load dual-task condition, and 65.63% (*SD* = 12.75) in the high load dual-task condition. The effect of task condition was significant [*F*(2,30) = 36.59, *p* < .001; η^2^_p_ = .71, BF = 6.75 × 10^6^]. Post hoc analysis revealed that auditory task accuracy was significantly lower in the high load dual-task condition compared to the other two conditions: vs. auditory single-task, *t* (15) = 7.38, *p* < .001, Cohen’s d = 1.84, BF = 8.31 × 10^3^; vs. low load dual-task, *t* (15) = 6.34, *p* < .001, Cohen’s d = 1.58, BF = 1.70 × 10^3^. The auditory task accuracy was not significantly different between auditory single-task and low load dual-task conditions [*t* (15) = 1.75, *p* = .10, Cohen’s d = 0.44, BF = 0.88].

Further, we examined the relationship between visual and auditory task performance during the dual-task conditions. The change in auditory accuracy [i.e., (low load – high load)/low load] was negatively correlated with the change in visual RT [i.e., (high load – low load)/low load] (Spearman’s ρ = −.46, uncorrected *p* = .038, one-sided; Figure 3D), such that listeners who slowed down more on the visual task from low to high load conditions tended to have a smaller drop in auditory accuracy. The change in auditory accuracy was not significantly correlated with the change in visual accuracy (Spearman’s ρ = −.29, uncorrected *p* = .28, one-sided).

These results demonstrate that divided (vs. selective) attention and increasing visual load impair behavioral visual and auditory performance.

### Neural tracking of visual stimuli is strongly modulated by divided attention and visual load

To assess neural tracking of visual stimuli, we focused on the predictive power of visual predictors while controlling for all speech-related predictors (acoustic and linguistic). Adding predictors for visual stimuli to a model including only speech predictors significantly improved its predictive power (prediction accuracy averaged across task conditions; *t*_max_ = 12.93, *p* < .001), providing evidence for neural tracking of visual stimuli. The cluster-based test resulted in a single significant cluster that spread across all electrodes, with the largest effects on parietal and occipital electrodes (Figure 4A).

**Figure 4.**
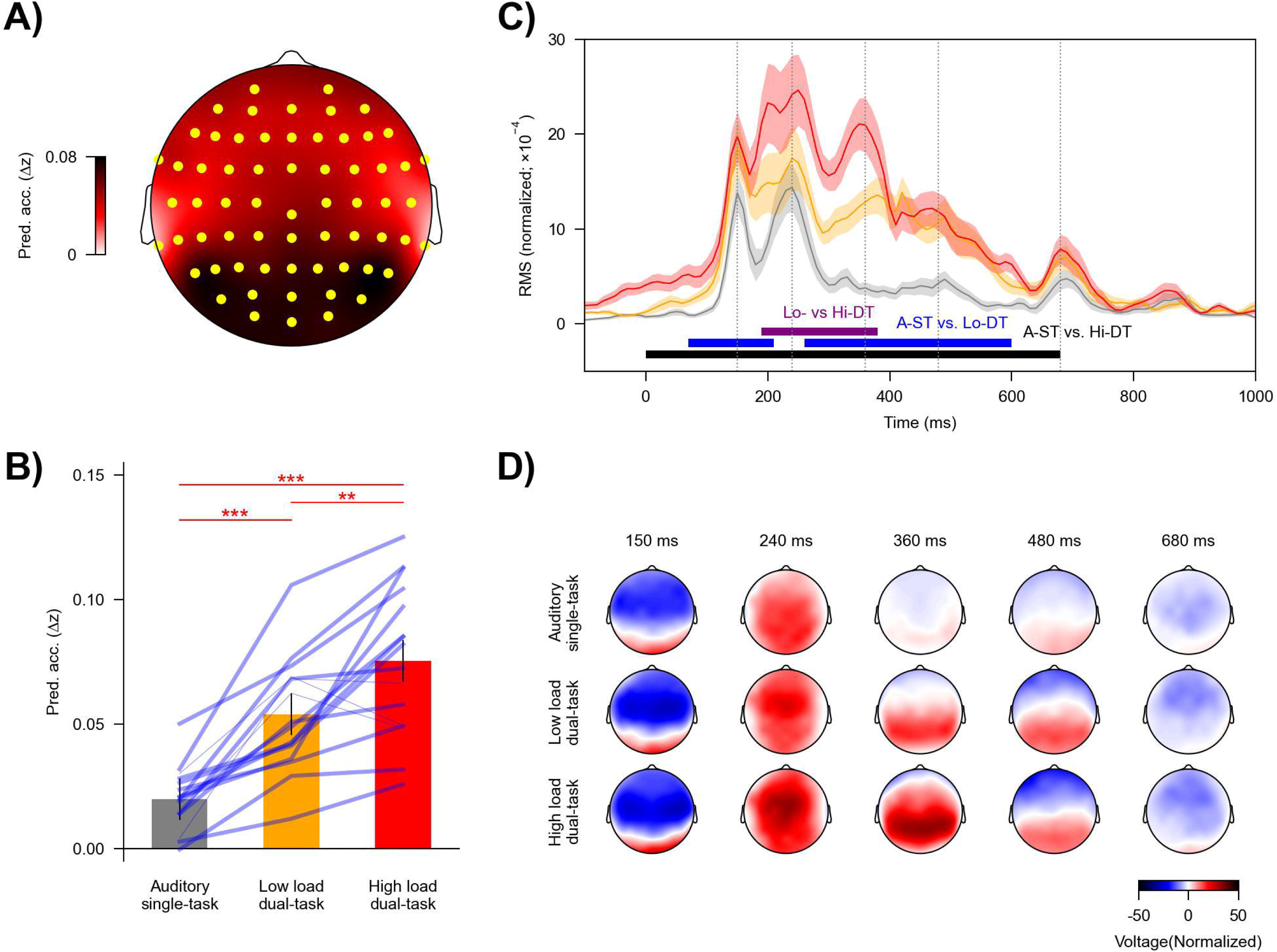
Neural tracking of visual stimuli across task conditions. Visual stimuli were associated with a robust response, which further increased with task-relevance and -load. (A) Topography showing the increase in prediction accuracy (Δz) due to visual predictors, which was significantly above zero in a single cluster encompassing the highlighted (yellow) electrodes. (B) Prediction accuracy across task conditions. Blue lines denote individual participants: Thicker lines indicate higher prediction accuracy for the high vs. low load condition, and thinner lines indicate lower accuracy for the high vs low load condition. Red asterisks denote *p* values for comparison between conditions. Error bars denote the 95% within-subject confidence interval (Loftus & Masson, 1994). (C) Global field power (GFP) of the visual temporal response functions (TRFs). Visual stimuli lasted from 0 to 500 ms. Shaded areas denote within-subject standard errors around the mean (for color labels see panel B). Horizontal lines denote time windows in which TRFs were significantly different between conditions. (D) Topographies of selected times in panel C (grey vertical lines). A-ST: auditory single-task, Lo-DT: low load dual-task, Hi-DT: high load dual-task. ** *p* < .01, *** *p* < .001.

Importantly, the predictive power of the visual predictors was modulated by task condition [*F*(2,30) = 46.10, *p* < .001, η^2^_p_ = .76, BH = 6.09 × 10^7^]. As shown in Figure 4B, the high load dual-task condition was associated with the highest predictive power (mean = 0.075, *SD* = 0.029), followed by the low load dual-task condition (mean = 0.053, *SD* = 0.022), and lowest for the auditory single-task condition (mean = 0.020, *SD* = 0.012): high load dual-task vs. auditory single-task, *t*(15) = 9.52, *p* < .001, Cohen’s d = 2.38, BF = 1.50 × 10^5^; high load dual-task vs. low load dual-task, *t*(15) = 3.34, *p* = .005, Cohen’s d = .84, BF = 10.7; low load dual-task vs. auditory single-task, *t*(15) = 6.64, *p* < .001, Cohen’s d = 1.66, BF = 2.72 × 10^3^. Together, these results suggest that neural tracking of visual stimuli was successively enhanced with increasing load of the visual task.

We analyzed mTRFs to further clarify how the difference in model predictive power was reflected in brain responses. Visual mTRFs can be conceptualized as evoked responses to the visual stimuli. Consistent with results for prediction accuracy, the mTRFs were also modulated by the task condition (Figure 4C). The high load dual-task condition showed larger mTRF amplitudes than the auditory single-task condition from 0 to 680 ms (*p* < .001) and the low load dual-task condition from 190 to 380 ms (*p* < .001). The mTRF amplitudes for the low load dual-task condition were larger than the auditory single-task condition from 70 to 210 ms (*p* = .009) and from 260 to 600 ms (*p* < .001).

### Divided attention, but not visual load, reduces late neural tracking of acoustic features

The acoustic predictors significantly contributed to model prediction beyond the visual and linguistic predictors in a cluster that spread across almost all electrodes, with maxima at temporal sites (*t*_max_ = 12.00, *p* < .001; Figure 5A). As expected, these results provide evidence for robust neural tracking of acoustic properties of speech.

**Figure 5.**
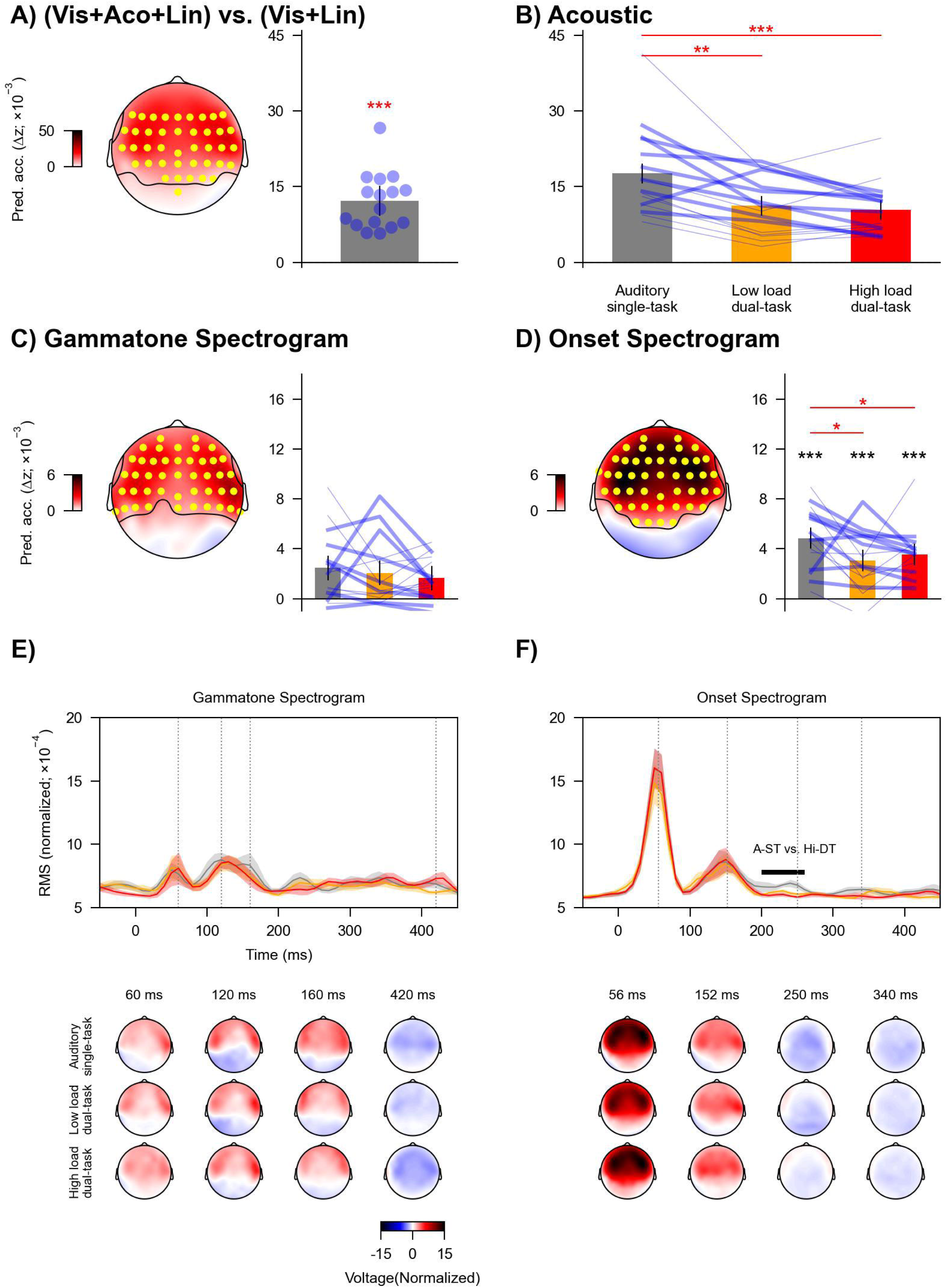
Neural tracking of acoustic information across task conditions. (A) Increase in prediction accuracy (Δz) due to acoustic predictors of speech (gammatone and onset spectrogram), which was significantly above zero in a cluster encompassing the highlighted (yellow) electrodes. Blue dots denote individual participants. (B) Prediction accuracy across task conditions for acoustic predictors, i.e., combined gammatone and onset spectrogram. (C) and (D) Prediction accuracy across task conditions for gammatone spectrogram and onset spectrogram separately. Topographies highlight electrodes (yellow) with prediction accuracy that was significantly above zero. Black asterisks denote *p* values for testing against (above) zero at individual conditions. (B) to (D) Blue lines denote individual participants: Thicker lines indicate lower accuracy for high vs. low load condition, and thinner lines indicate higher accuracy for high vs. low load condition. Red asterisks denote *p* values for comparison between conditions. Error bars denote 95% confidence interval. (E) and (F) Global field power (GFP; top) of mTRFs and related topographies (bottom) for gammatone and onset spectrogram. The mTRFs were averaged across the eight frequency bands. Shaded areas denote within-subject standard errors around the mean. Horizontal lines denote time windows in which mTRFs were significantly different between conditions. Topographies are shown for selected times indicated in GFPs (grey vertical lines). A-ST: auditory single-task, Lo-DT: low load dual-task, Hi-DT: high load dual-task. * *p* < .05, ** *p* < .01, *** *p* < .001.

The prediction accuracy for acoustic predictors was modulated by task condition [*F*(2,30) = 14.83, *p* < .001, η^2^_p_ = .50, BF = 581.38; Figure 5B]. Post hoc analysis showed that the prediction accuracy significantly dropped in the two dual-task conditions compared to the auditory single-task condition [vs. low load dual-task, *t*(15) = 3.84, *p* = .002, Cohen’s d = 0.96, BF = 25.60; vs. high load dual-task, *t*(15) = 4.78, *p* < .001, Cohen’s d = 1.20, BF = 130.60]. The prediction accuracy was not significantly different between the dual-task conditions [*t*(15) = .77, *p* = .45, Cohen’s d = 0.19, BF = 0.33]. These results suggest that neural tracking of acoustic information was reduced when directing attention from one task (auditory) to two tasks (visual and auditory).

Then, we assessed whether the effect of task condition could be attributed to specific acoustic predictors. The two acoustic predictors both independently contributed to overall model prediction (gammatone spectrogram: *t*_max_ = 6.08, *p* < .001, Figure 5C; onset spectrogram: *t*_max_ = 9.91, *p* < .001, Figure 5D). The effect of task condition on the prediction accuracy was significant for onset spectrogram [*F*(2,30) = 4.93, *p* = .033, η^2^_p_ = 0.25, BF = 4.17] but not for gammatone spectrogram [*F*(2,30) = .70, *p* = .59, η^2^_p_ = 0.04, BF = 0.26]. Post hoc analysis revealed that the prediction accuracy for onset spectrogram significantly dropped in the two dual-task conditions compared to the auditory single-task condition [vs. low load dual-task, *t*(15) = 2.61, *p* = .030, Cohen’s d = 0.65, BF = 3.14; vs. high load dual-task, *t*(15) = 2.89, *p* = .030, Cohen’s d = 0.72, BF = 4.94]. The prediction accuracy was not significantly different between the dual-task conditions [*t*(15) = −.79, *p* = .44, Cohen’s d = 0.20, BF = 0.34].

Considering the modulation by task condition, we further examined whether divided attention eliminated neural tracking of onset spectrogram. The prediction accuracy at individual task conditions was significantly above zero (all uncorrected *p*s < .001, Cohen’s d > 1.20, BF > 256.40; Figure 5D), suggesting that directing attention from one task to two tasks did not eliminate the neural tracking of acoustic onsets.

Finally, we examined the effect of task condition on the mTRFs for the onset spectrogram (Figure 5F). mTRFs to a continuous stimulus like the auditory spectrogram can be conceived of as evoked responses to an elementary event in the stimulus (i.e., the impulse response). The mTRF amplitudes in the auditory single-task condition were larger compared to the high load dual-task condition from 200 to 260 ms (*p* = .003). Further, a visual inspection of the mTRFs from individual subjects revealed two relatively reliable peaks at about 56 (P1) and 152 (P2) ms. Latencies of these peaks were not significantly different across conditions [56 ms: *F*(2,30) = .65, uncorrected *p* = .94, η^2^_p_ = 0.004; 152 ms: *F*(2,30) = .62, uncorrected *p* = .54, η^2^_p_ = 0.04].

In sum, acoustic tracking was very similar across conditions, with only a slight reduction in the tracking of acoustic onsets in the divided attention tasks, compared to the single task. This difference was likely explained by a reduction in a relatively late response component, starting at 200 ms.

### Divided attention, but not visual load, reduces late tracking of linguistic information

The linguistic predictors significantly contributed to model prediction beyond the visual and acoustic predictors (*t*_max_ = 4.95, *p* < .001; Figure 6A). The cluster-based test indicated that the effect of linguistic predictors was primarily observed for temporal-central electrodes. These results provide evidence for neural tracking of linguistic properties of speech.

**Figure 6.**
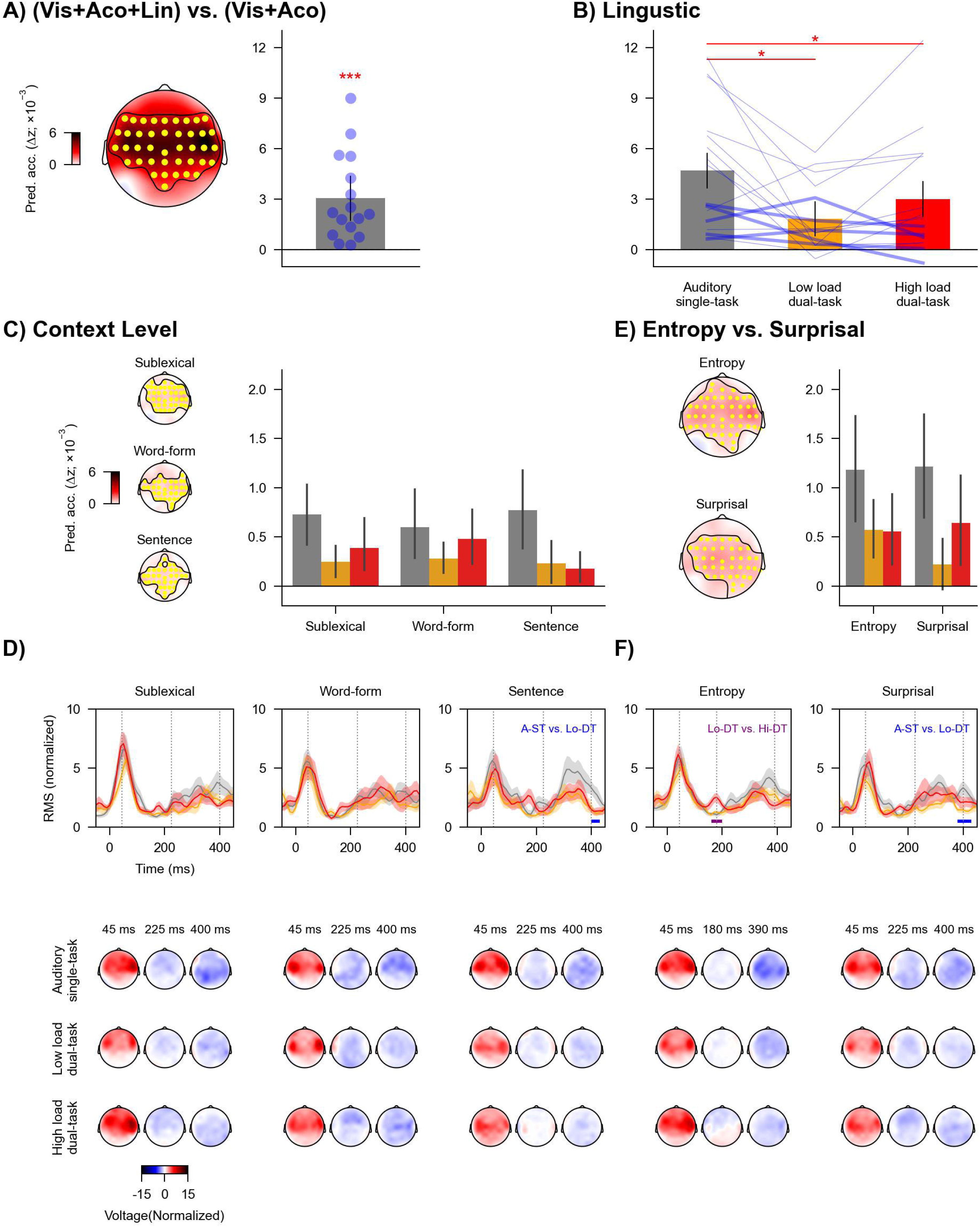
Neural tracking of linguistic information across task conditions. (A) Increase in prediction accuracy (Δz) due to linguistic predictors of speech (word onsets, phoneme onsets, sublexical surprisal and entropy, word-form surprisal and entropy, and sentence surprisal and entropy), which was significantly above zero across highlighted (yellow) electrodes in the topography. Blue dots denote individual participants. (B) Prediction accuracy for combined linguistic predictors across conditions. Blue lines denote individual participants: Thicker lines indicate lower accuracy for high vs. low load condition, and thinner lines indicate higher accuracy for high vs. low load condition. Red asterisks denote *p* values for comparison between conditions. (C) Prediction accuracy for three context levels (sublexical, word-form, and sentence) across conditions. Each level includes entropy and surprisal predictors. (D) Global field power (GFP; top) of mTRFs and related topographies (bottom) for each context level. The mTRF GFPs were averaged across entropy and surprisal. (E) Prediction accuracy for entropy and surprisal. Each predictor includes the three context levels. (F) GFP of mTRFs and related topographies for entropy and surprisal. (B), (C), and (E) Error bars denote 95% confidence interval. (D) and (F) Shaded areas denote standard errors around the mean. Horizontal lines denote time windows in which the mTRFs were significantly different between task conditions. Topographies are shown for selected times indicated in GFPs (grey vertical lines). A-ST: auditory single-task, Lo-DT: low load dual-task, Hi-DT: high load dual-task. * *p* < .05, *** *p* < .001.

The prediction accuracy for linguistic predictors was modulated by task condition [*F*(1.41, 21.15) = 6.66, *p* = .029, η^2^_p_ = 0.31, BF = 10.82; Figure 6B]. The prediction accuracy significantly dropped in the two dual-task conditions compared to the auditory single-task condition [vs. low load dual-task, *t*(15) = 2.83, *p* = .029, Cohen’s d = 0.71, BF = 4.49; vs. high load dual-task, *t*(15) = 2.61, *p* = .029, Cohen’s d = 0.65, BF = 3.16]. The prediction accuracy was not significantly different between the two dual-task conditions [*t*(15) = −1.80, *p* = .091, Cohen’s d = 0.45, BF = 0.95]. These results suggest that neural tracking of linguistic information was reduced when directing attention from one task to two tasks.

Next, we conducted three sets of analyses to assess whether the effect of task condition could be attributed to specific linguistic properties.

#### Task effects appear to be similar across different context levels

The first analysis focused on the three context levels (sublexical, word-form, and sentence). Each level independently contributed significantly to model prediction (sublexical: *t*_max_ = 5.22, *p* < .001; word-form: *t*_max_ = 3.92, *p* < .001; sentence: *t*_max_ = 4.98, *p* < .001; Figure 6C). A two-way repeated-measures ANOVA showed that the interaction between context level and task condition was not significant [*F*(2.55, 38.18) = 1.19, *p* = .40, η^2^_p_ = 0.073, BF = 0.19]. The main effect of context level was not significant [*F*(2,30) = .32, *p* = .77, η^2^_p_ = 0.021, BF = 0.08]. But the main effect of task condition was significant [*F*(1.2,18.01) = 8.46, *p* = .021, η^2^_p_ = 0.36, BF = 1.40 × 10^3^]. Post hoc analysis showed that the prediction accuracy was significantly reduced from the auditory single-task condition to the low load [*t*(15) = 2.90, *p* = .016, Cohen’s d = 0.73, BF = 5.08] and high load dual-task conditions [*t*(15) = 4.27, *p* = .002, Cohen’s d = 1.07, BF = 54.63]. But the prediction accuracy was not significantly different between the low and high load dual-task condition[*t*(15) = 1.02, *p* = .32, Cohen’s d = 0.26, BF = 0.40]. Further, we found similar patterns of results when restricting the two-way repeated-measures ANOVA analysis to the dual-task conditions. In sum, patterns of task condition effects observed for linguistic predictors appeared to be similar across the different linguistic models.

Considering the modulation by context level and task condition, we further examined whether divided attention eliminated neural tracking of any of these predictors. The prediction accuracies for all predictors at individual task conditions were significantly above zero (all uncorrected *p*s < .03, Cohen’s d > 0.53, BF > 1.51).

Regarding mTRFs, the effect of task condition was not significant for sublexical or word-form context but was for sentence context (Figure 6D). The mTRF amplitude of sentence context in the auditory single-task condition was larger compared to the low load dual-task condition from 400 to 430 ms (*p* = .036). Topographies suggest that this is due to a broadly distributed more negative component in the single task condition.

Initial response peaks to linguistic features appear relatively early. This is consistent with previous results (Brodbeck et al., 2022) and might be partly because forced alignment, which was used to determine phoneme timing, does not take into account coarticulation effects. Some information about upcoming phonetic features might thus have systematically precede the estimates of phoneme onset times we used.

#### Neural tracking of surprisal might increase with visual load

The second analysis focused on entropy and surprisal. The two predictors independently contributed significantly to model prediction (entropy: *t*_max_ = 5.51, *p* < .001; surprisal: *t*_max_ = 3.91, *p* = .001; Figure 6E). A two-way repeated-measures ANOVA showed that the interaction between predictor type (entropy vs. surprisal) and task condition was not significant [*F*(1.32, 19.86) = 1.29, *p* = .40, η^2^_p_ = 0.079, BF = 0.31]. The main effect of predictor type was not significant [*F*(1,15) = .31, *p* = .65, η^2^_p_ = 0.02, BF = 0.23]. But the main effect of task condition was significant [*F*(1.35,20.2) = 9.85, *p* = .011, η^2^_p_ = 0.40, BF = 890.10]. Post hoc analysis showed that, when averaging across surprisal and entropy, the prediction accuracy was significantly reduced from the auditory single-task condition to the low load [*t*(15) = 3.28, *p* = .008, Cohen’s d = 0.82, BF = 9.56] and high load dual-task conditions [*t*(15) = 4.11, *p* = .003, Cohen’s d = 1.03, BF = 40.91]. Numerically, the prediction accuracy was improved from the low load to high load dual-task condition, but this difference was not significant [*t*(15) = 1.27, *p* = .22, Cohen’s d = 0.32, BF = 0.51].

Because of theoretical predictions of enhanced reliance on linguistic representations during higher visual task load (see *Introduction* and *Discussion*), we further restricted the two-way repeated-measures ANOVA analysis to the dual-task conditions. The interaction between predictor type and task condition was significant [*F*(1,15) = 5.75, uncorrected *p* = .03 (FDR-corrected *p* = .063), η^2^_p_ = 0.28, BF = 1.23]. There was no significant main effect of predictor type [*F*(1,15) = 1.92, *p* = .33, η^2^_p_ = 0.11, BF = 0.40] or task condition [*F*(1,15) = 1.62, *p* = .36, η^2^_p_ = 0.097, BF = 0.79]. Post hoc analysis showed that for entropy, the prediction accuracy was not different between the dual-task conditions [*t*(15) = .10, *p* = .92, Cohen’s d = 0.03, BF = 0.26]. But for surprisal, the prediction accuracy was significantly improved from the low load to high load dual-task condition [*t*(15) = 2.20, uncorrected *p* = .044, Cohen’s d = 0.55, BH = 1.66].

Considering the modulation by predictor type and task condition, we further examined whether divided attention eliminated neural tracking of entropy or surprisal. The prediction accuracies for both predictors at individual task conditions were significantly above zero (all uncorrected *p*s < .01, Cohen’s d > 0.66, BF > 3.36), except for the surprisal predictors at the low load dual-task condition (uncorrected *p* = .059, Cohen’s d = 0.41, BF = 0.78).

Regarding mTRFs, the mTRF amplitude of entropy in the low load dual-task condition was smaller than the high load dual-task condition from 160 to 200 ms (uncorrected *p* = .037). The mTRF amplitude of surprisal in the low load dual-task condition was smaller compared to the auditory single-task condition from 380 to 430 ms (uncorrected *p* = .012). We did not observe a significant effect of task load, although the mTRF to surprisal during high visual load was numerically stronger than low load from 200 ms onwards.

### Divided attention or visual load does not affect neural tracking of word onsets

The third set of analysis focused on word onset. This predictor independently contributed significantly to model prediction (*t*_max_ = 4.51, *p* < .001; Figure 7A). But the effect of task condition on prediction accuracy was not significant [*F*(2,30) = .07, *p* = .93, η^2^_p_ = 0.005, BF = 0.17; Figure 7B].

**Figure 7.**
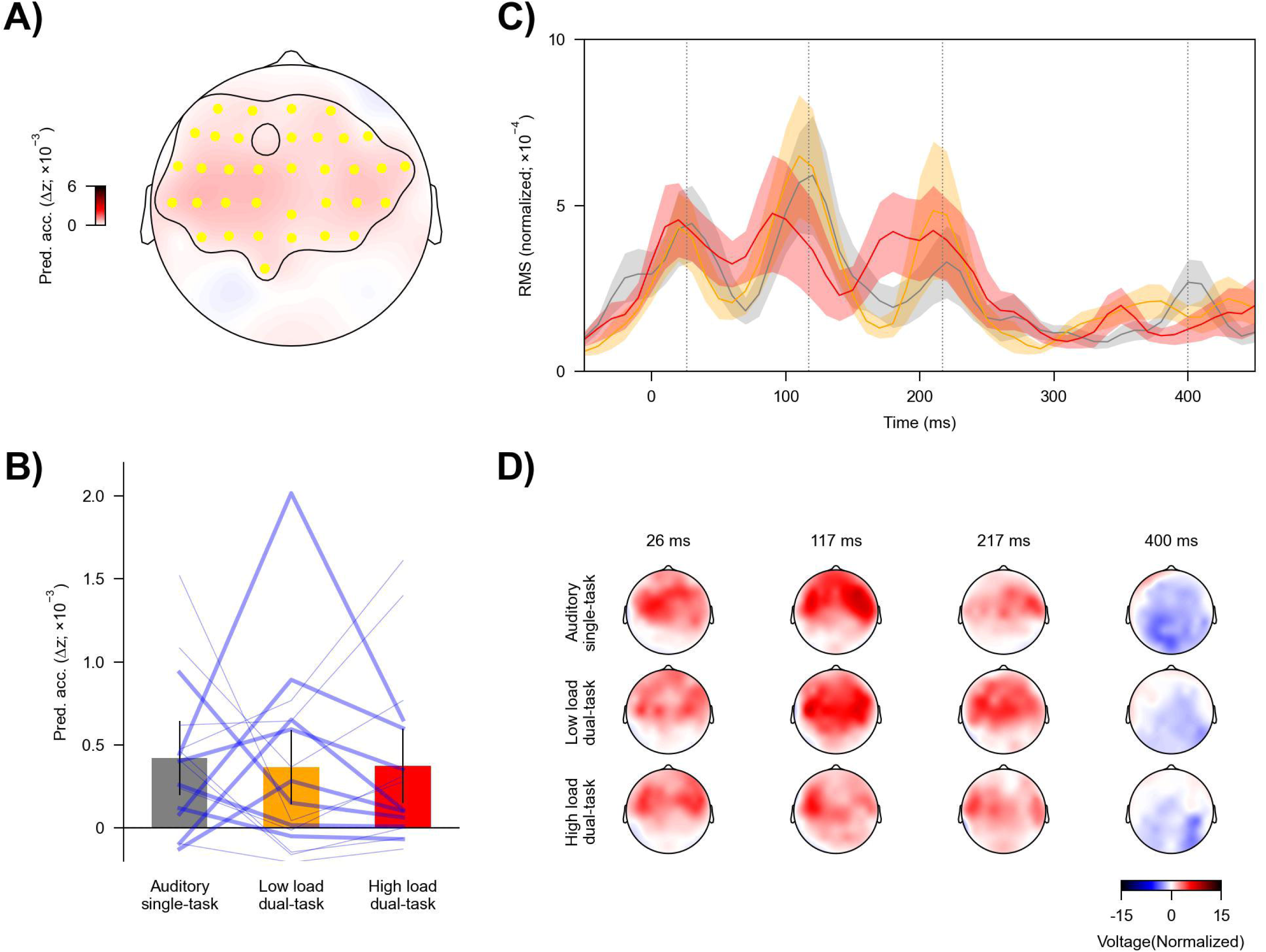
Neural tracking of word onsets across task conditions. (A) Topography showing the increase in prediction accuracy (Δz) due to word onsets, which was significantly above zero across highlighted (yellow) electrodes. (B) Prediction accuracy across conditions. Blue lines denote individual participants: Thicker lines indicate higher accuracy for the high vs. low load condition, and thinner lines indicate lower accuracy for the high vs low load condition. Error bars denote 95% within-subject confidence interval (Loftus & Masson, 1994). (C) Global field power (GFP) of mTRFs. Shaded areas denote within-subject standard error around the mean. (D) Topographies of selected times in panel C (grey vertical lines). * *p* < .05, ** *p* < .01.

Taken together, results suggest that directing attention from one task to two tasks may reduce but does not eliminate the neural tracking of linguistic features of speech. However, increasing visual load does not lead to a further reduction. On the contrary, an increasing load of the dual task might even be associated with enhanced neural tracking of phoneme surprisal. However, this effect should be interpreted with care because the effect was not significant when analyzing all linguistic predictors as a group or after correction for multiple comparisons.

## Discussion

We examined the extent to which bimodal divided attention influences acoustic and linguistic representations of natural continuous speech. Compared to unimodal auditory speech processing, the visual tasks affected acoustic onsets (but not the acoustic spectrogram, Figure 5D) and linguistic representations related to predictive processing (but not to lexical segmentation, Figure 6E). Surprisingly, we did not find evidence of further reduction (at any processing level) with increased visual (dual) task load, despite unambiguous behavioral and neural evidence of the high load task as being more demanding (Figures 3 and 4).

### Locus of effects of bimodal divided attention on continuous speech processing

We noted a striking dissociation between the impact of the dual-task on behavioral performance in the speech comprehension task, and a relative lack of impact on neural speech processing. Behaviorally, the dual-task load clearly impacted listeners’ ability to answer auditory comprehension questions. However, neural tracking of acoustic and linguistic speech features was affected only at late response components, and remained largely unchanged with varying dual-task load. This neural and behavioral dissociation suggests that bimodal divided attention largely only impacts late, post-perceptual processes of continuous speech processing. The significant and unchanged responses related to predictive processing using the sentence context suggest that listeners could track multi-word sequences regardless of dual-task load. We posit that the decreased behavioral performance originates from higher-order cognitive processes that are not adequately described by probabilistic word-sequence models, such as semantic integration and memory formation.

Previous behavioral research has suggested that increased dual-task load is associated with reduced acoustic sensitivity during speech recognition (Mattys et al., 2014; Mattys & Palmer, 2015; Mattys & Wiget, 2011). In our data, the dual-task did not alter early acoustic responses and only had subtle effects on later (> 200 ms) responses (see Figures 5E and 5F). This suggests that the effect of bimodal divided attention may not be on basic acoustic representations per se, but on secondary acoustic analysis stages or on how these representations are accessed and used by higher-order processes.

### Implications for resource-based accounts

A common framework for understanding effects under dual-task paradigms is resource-based (Wahn & König, 2017). When two tasks draw from a limited pool of shared resources, increased load in one task is associated with poorer performance in another task. Such a hypothesis is often referred to as the *supramodal* account of crossmodal attention (Broadbent, 1958; Ciaramitaro et al., 2017; Klemen et al., 2009; Macdonald & Lavie, 2011; Molloy et al., 2015). In contrast, if the increased load in one task does not affect a corresponding decrease in another, the two modalities can draw on separate resource pools. Such a hypothesis is consistent with a *modality-specific* account of crossmodal attention (Alais et al., 2006; Arrighi et al., 2011; Duncan et al., 1997; Keitel et al., 2013; Parks et al., 2011; Porcu et al., 2014).

Here, we observed a reduction in neural tracking of speech acoustic and linguistic features under bimodal divided attention, consistent with previous studies demonstrating detrimental effects of bimodal divided attention for simplified speech stimuli such as syllables (Gennari et al., 2018), words (Kasper et al., 2014), and sentences (Salo et al., 2015). In conjunction with the co-occurring improved neural tracking of visual stimuli, this finding appears to suggest a tradeoff between attending to the auditory versus visual modalities. Hence, these results appear to align with the *supramodal* hypothesis of the dual-task effects that the auditory and visual tasks of our study draw on a limited pool of shared resources (Broadbent, 1958; Ciaramitaro et al., 2017; Klemen et al., 2009; Macdonald & Lavie, 2011; Molloy et al., 2015).

However, a *supramodal* hypothesis of the dual-task effects does not seem to fit other key results from our study. First, the impact of bimodal divided attention is specific to certain features of the speech signals: we found bimodal attention effects for acoustic onsets but not acoustic envelope representations, and for predictive linguistic processing, indexed through information-theoretic variables, but not lexical segmentation, indexed through the word-onset predictors. In each case, the impact of divided attention is not a generally reduced representation but is restricted to only specific response components in the response time courses (the mTRFs).

Furthermore, a resource-based account would suggest that when the visual load is further increased, available resources for speech representations should further decrease, which is not what we observed. Instead, adding a visual task exacted a cost on neural speech representations, but this cost did not scale with the task load. In contrast to these neural effects, task load did affect behavioral performance on the auditory task. These divergent results may require an explanation involving different resource pools (Wahn & König, 2017). For example, there may be a resource pool for sensory processing, which is sensitive to divided attention but not task load, thus, is relatively modality-specific. There may be a second resource pool, which is sensitive to task load and affects higher-order story comprehension, thus, is relatively supramodal.

### Selective *versus* divided attention on speech processing

Previous studies on continuous speech processing have shown that selective attention within and between modalities strongly modulates neural processing of both acoustic and linguistic features of continuous speech, and the attentional effects seem to be even stronger for linguistic processing (Brodbeck et al., 2018; Broderick et al., 2018; Ding et al., 2018; Kiremitçi et al., 2021; Vanthornhout et al., 2019; Yahav & Golumbic, 2021). Specifically, neural tracking of acoustic features is reduced and delayed but not eliminated for unattended speech, but the tracking of linguistic features is virtually abolished. A parsimonious null hypothesis, consistent with the notion of a shared resource pool, is that speech representations during divided attention ought to be halfway between attended and ignored speech. Our results suggest that this is not the case.

First, certain speech features (e.g., acoustic spectrogram and word onsets) that have been shown in prior work to be modulated by selective attention are insensitive to bimodal divided attention. Second, unlike prior work demonstrating differential selective attention effects on the relative balance of acoustic vs. linguistic processing, we did not observe a greater reduction in linguistic processing than acoustic processing with the manipulation of divided attention. The neural tracking of both feature classes is reduced but not eliminated. Third, for those features showing modulation by divided attention, we did not observe any delay in the neural responses as reflected in the mTRFs (Figures 5 and 6). Fourth, the effect of divided attention emerged largely at later stages (after ~200 ms) with the earlier latencies relatively unaffected. Thus, the effect of bimodal divided attention on neural continuous speech processing appears to be feature-specific and occurs relatively late in processing.

These differences indicate that selective and divided attention are subserved by distinct mechanisms. Relative to selective attention, bimodal divided attention tasks may be associated with additional recruitment of frontal regions that interact with sensory cortices (Gennari et al., 2018; Johnson & Zatorre, 2006; Loose et al., 2003). A stronger engagement of frontal regions has been associated with poorer task performance (Gennari et al., 2018; Johnson & Zatorre, 2006). These neural findings appear to align with the argument that the costs of bimodal divided attention may come from limitations of executive control to coordinate processes related to two tasks rather than a competition for shared sensory resources (Katus & Eimer, 2019; Loose et al., 2003). The differential effects of selective and divided attention on continuous speech processing suggest that the costs of selective attention are more likely to originate from ‘filter’ mechanisms (Broadbent, 1958; Lachter et al., 2004) that pass task-relevant signals but block task-irrelevant others, instead of the re-allocation of shared resources. Nevertheless, future studies are needed to elucidate mechanisms underlying differences in continuous speech processing between selective and divided attention.

### Increased responses to surprisal with dual-task load

We found that increasing visual load increased responses to phoneme surprisal, but not entropy. This effect was statistically only seen after excluding the auditory single-task condition and should thus be interpreted with care, but it is consistent with several extant findings. The dissociation between entropy and surprisal is consistent with recent evidence that these two processes may reflect different neural processes (Gaston et al., 2022). Neural responses associated with surprisal may reflect prediction errors that signal the difference between predicted and observed phonemes. Such prediction error signals may be boosted when attention is directed to the speech stimuli (e.g., auditory single-task; Auksztulewicz & Friston, 2015; Smout et al., 2019) or when attention to the speech stimuli is directed away to demanding crossmodal tasks (e.g., high-load visual tasks; Xie et al., 2018). The increased response to surprisal might also reflect a shift toward more reliance on linguistic representations during speech processing when resources for auditory processing were constrained under divided attention of higher load (Mattys et al., 2009; Mattys & Wiget, 2011).

### Neural tracking of word onsets was not affected by divided attention

Tracking of word onsets might reflect lexical segmentation (Sanders et al., 2002; Sanders & Neville, 2003) and, along with other linguistic features, is strongly affected by selective attention (Brodbeck et al., 2018). It has been suggested that neural responses to word onsets reflect the dynamic allocation of attention to time windows that contain word onsets (Astheimer & Sanders, 2009). However, our results indicate that tracking of word onsets is robust to manipulations of attentional load by adding a visual task and increasing dual-task load. This suggests that the word-onset attention effect may draw on a relatively unshared resource pool, or that the word-onset responses reflect a more mechanistic aspect of lexical segmentation.

## Conclusion

This study demonstrates a striking dissociation between the impact of dual-task load on behavioral speech comprehension performance and a relative lack of impact on time-locked neural representations of continuous speech. The behavioral effects of bimodal divided attention on continuous speech processing occur not due to impaired early sensory representations but likely at later cognitive processing stages.

## Acknowledgments

We thank Rachel Reetzke, Elise LeBovidge, and Jacie McHaney for their assistance with participant recruitment and data collection. The content is solely the responsibility of the authors and does not necessarily represent the official views of the National Institutes of Health or the Natural Science Foundation.

## References

Alais, D., Morrone, C., & Burr, D. (2006). Separate attentional resources for vision and audition. Proceedings of the Royal Society B: Biological Sciences, 273(1592), 1339–1345.

Arrighi, R., Lunardi, R., & Burr, D. (2011). Vision and audition do not share attentional resources in sustained tasks. Frontiers in Psychology, 2, 56.

Astheimer, L. B., & Sanders, L. D. (2009). Listeners modulate temporally selective attention during natural speech processing. Biological Psychology, 80(1), 23–34.

Auksztulewicz, R., & Friston, K. (2015). Attentional enhancement of auditory mismatch responses: A DCM/MEG study. Cerebral Cortex, 25(11), 4273–4283.

Benjamini, Y., & Hochberg, Y. (1995). Controlling the false discovery rate: A practical and powerful approach to multiple testing. Journal of the Royal Statistical Society: Series B (Methodological), 57(1), 289–300.

Bidelman, G. M., & Alain, C. (2015). Musical training orchestrates coordinated neuroplasticity in auditory brainstem and cortex to counteract age-related declines in categorical vowel perception. Journal of Neuroscience, 35(3), 1240–1249.

Broadbent, D. E. (1958). Perception and communication. Pergamon Press.

Brodbeck, C., Bhattasali, S., Heredia, A. A. C., Resnik, P., Simon, J. Z., & Lau, E. (2022). Parallel processing in speech perception with local and global representations of linguistic context. Elife, 11, e72056.

Brodbeck, C., Das, P., Kulasingham, J. P., Bhattasali, S., Gaston, P., Resnik, P., & Simon, J. Z. (2021). Eelbrain: A Python toolkit for time-continuous analysis with temporal response functions. BioRxiv.

Brodbeck, C., Hong, L. E., & Simon, J. Z. (2018). Rapid transformation from auditory to linguistic representations of continuous speech. Current Biology, 28(24), 3976–3983.

Brodbeck, C., Jiao, A., Hong, L. E., & Simon, J. Z. (2020). Neural speech restoration at the cocktail party: Auditory cortex recovers masked speech of both attended and ignored speakers. PLoS Biology, 18(10), e3000883.

Brodbeck, C., & Simon, J. Z. (2020). Continuous speech processing. Current Opinion in Physiology, 18, 25–31.

Broderick, M. P., Anderson, A. J., Di Liberto, G. M., Crosse, M. J., & Lalor, E. C. (2018). Electrophysiological correlates of semantic dissimilarity reflect the comprehension of natural, narrative speech. Current Biology, 28(5), 803–809.

Ciaramitaro, V. M., Chow, H. M., & Eglington, L. G. (2017). Cross-modal attention influences auditory contrast sensitivity: Decreasing visual load improves auditory thresholds for amplitude-and frequency-modulated sounds. Journal of Vision, 17(3), 20–20.

Crosse, M. J., Di Liberto, G. M., Bednar, A., & Lalor, E. C. (2016). The multivariate temporal response function (mTRF) toolbox: A MATLAB toolbox for relating neural signals to continuous stimuli. Frontiers in Human Neuroscience, 10, 604.

Ding, N., Pan, X., Luo, C., Su, N., Zhang, W., & Zhang, J. (2018). Attention is required for knowledge-based sequential grouping: Insights from the integration of syllables into words. Journal of Neuroscience, 38(5), 1178–1188.

Ding, N., & Simon, J. Z. (2012). Neural coding of continuous speech in auditory cortex during monaural and dichotic listening. Journal of Neurophysiology, 107(1), 78–89.

Duncan, J., Martens, S., & Ward, R. (1997). Restricted attentional capacity within but not between sensory modalities. Nature, 387(6635), 808–810.

Fishbach, A., Nelken, I., & Yeshurun, Y. (2001). Auditory edge detection: A neural model for physiological and psychoacoustical responses to amplitude transients. Journal of Neurophysiology, 85(6), 2303–2323.

Gaston, P., Brodbeck, C., Phillips, C., & Lau, E. (2022). Auditory Word Comprehension is Less Incremental in Isolated Words. Neurobiology of Language, 1–50. https://doi.org/10.1162/nol_a_00084

Gennari, S. P., Millman, R. E., Hymers, M., & Mattys, S. L. (2018). Anterior paracingulate and cingulate cortex mediates the effects of cognitive load on speech sound discrimination. NeuroImage, 178, 735–743.

Gillis, M., Van Canneyt, J., Francart, T., & Vanthornhout, J. (2022). Neural tracking as a diagnostic tool to assess the auditory pathway. Hearing Research, 108607.

Gramfort, A., Luessi, M., Larson, E., Engemann, D. A., Strohmeier, D., Brodbeck, C., Goj, R., Jas, M., Brooks, T., & Parkkonen, L. (2013). MEG and EEG data analysis with MNE-Python. Frontiers in Neuroscience, 267.

Hamilton, L. S., & Huth, A. G. (2020). The revolution will not be controlled: Natural stimuli in speech neuroscience. Language, Cognition and Neuroscience, 35(5), 573–582.

Heafield, K. (2011). KenLM: Faster and smaller language model queries. 187–197.

Hickok, G., & Poeppel, D. (2007). The cortical organization of speech processing. Nature Reviews Neuroscience, 8(5), 393–402.

Jaeggi, S. M., Buschkuehl, M., Etienne, A., Ozdoba, C., Perrig, W. J., & Nirkko, A. C. (2007). On how high performers keep cool brains in situations of cognitive overload. Cognitive, Affective, & Behavioral Neuroscience, 7(2), 75–89.

Johnson, J. A., & Zatorre, R. J. (2006). Neural substrates for dividing and focusing attention between simultaneous auditory and visual events. Neuroimage, 31(4), 1673–1681.

Kasper, R. W., Cecotti, H., Touryan, J., Eckstein, M. P., & Giesbrecht, B. (2014). Isolating the neural mechanisms of interference during continuous multisensory dual-task performance. Journal of Cognitive Neuroscience, 26(3), 476–489.

Katus, T., & Eimer, M. (2019). The sources of dual-task costs in multisensory working memory tasks. Journal of Cognitive Neuroscience, 31(2), 175–185.

Keitel, C., Maess, B., Schröger, E., & Müller, M. M. (2013). Early visual and auditory processing rely on modality-specific attentional resources. Neuroimage, 70, 240–249.

Keuleers, E., Brysbaert, M., & New, B. (2010). SUBTLEX-NL: A new measure for Dutch word frequency based on film subtitles. Behavior Research Methods, 42(3), 643–650.

Kiremitçi, I., Yilmaz, Ö., Çelik, E., Shahdloo, M., Huth, A. G., & Çukur, T. (2021). Attentional modulation of hierarchical speech representations in a multitalker environment. Cerebral Cortex, 31(11), 4986–5005.

Klemen, J., Büchel, C., & Rose, M. (2009). Perceptual load interacts with stimulus processing across sensory modalities. European Journal of Neuroscience, 29(12), 2426–2434.

Lachter, J., Forster, K. I., & Ruthruff, E. (2004). Forty-five years after Broadbent (1958): Still no identification without attention. Psychological Review, 111(4), 880.

Loftus, G. R., & Masson, M. E. (1994). Using confidence intervals in within-subject designs. Psychonomic Bulletin & Review, 1(4), 476–490.

Loose, R., Kaufmann, C., Auer, D. P., & Lange, K. W. (2003). Human prefrontal and sensory cortical activity during divided attention tasks. Human Brain Mapping, 18(4), 249–259.

Macdonald, J. S., & Lavie, N. (2011). Visual perceptual load induces inattentional deafness. Attention, Perception, & Psychophysics, 73(6), 1780–1789.

Maris, E., & Oostenveld, R. (2007). Nonparametric statistical testing of EEG-and MEG-data. Journal of Neuroscience Methods, 164(1), 177–190.

Marslen-Wilson, W. D. (1987). Functional parallelism in spoken word-recognition. Cognition, 25(1–2), 71–102.

Mattys, S. L., Barden, K., & Samuel, A. G. (2014). Extrinsic cognitive load impairs low-level speech perception. Psychonomic Bulletin & Review, 21(3), 748–754.

Mattys, S. L., Brooks, J., & Cooke, M. (2009). Recognizing speech under a processing load: Dissociating energetic from informational factors. Cognitive Psychology, 59(3), 203–243.

Mattys, S. L., & Palmer, S. D. (2015). Divided attention disrupts perceptual encoding during speech recognition. The Journal of the Acoustical Society of America, 137(3), 1464–1472.

Mattys, S. L., & Wiget, L. (2011). Effects of cognitive load on speech recognition. Journal of Memory and Language, 65(2), 145–160.

McAuliffe, M., Socolof, M., Mihuc, S., Wagner, M., & Sonderegger, M. (2017). Montreal Forced Aligner: Trainable Text-Speech Alignment Using Kaldi. 2017, 498–502.

Molloy, K., Griffiths, T. D., Chait, M., & Lavie, N. (2015). Inattentional deafness: Visual load leads to time-specific suppression of auditory evoked responses. Journal of Neuroscience, 35(49), 16046–16054.

Morey, R. D., Rouder, J. N., Jamil, T., & Morey, M. R. D. (2022). *BayesFactor: Computation of Bayes Factors for Common Designs*. https://CRAN.R-project.org/package=BayesFactor

Oostenveld, R., & Praamstra, P. (2001). The five percent electrode system for high-resolution EEG and ERP measurements. Clinical Neurophysiology, 112(4), 713–719.

Parks, N. A., Hilimire, M. R., & Corballis, P. M. (2011). Steady-state signatures of visual perceptual load, multimodal distractor filtering, and neural competition. Journal of Cognitive Neuroscience, 23(5), 1113–1124.

Pickering, M. J., & Gambi, C. (2018). Predicting while comprehending language: A theory and review. Psychological Bulletin, 144(10), 1002.

Porcu, E., Keitel, C., & Müller, M. M. (2014). Visual, auditory and tactile stimuli compete for early sensory processing capacities within but not between senses. Neuroimage, 97, 224– 235.

Salo, E., Rinne, T., Salonen, O., & Alho, K. (2015). Brain activations during bimodal dual tasks depend on the nature and combination of component tasks. Frontiers in Human Neuroscience, 9, 102.

Sanders, L. D., & Neville, H. J. (2003). An ERP study of continuous speech processing: I. Segmentation, semantics, and syntax in native speakers. Cognitive Brain Research, 15(3), 228–240.

Sanders, L. D., Newport, E. L., & Neville, H. J. (2002). Segmenting nonsense: An event-related potential index of perceived onsets in continuous speech. Nature Neuroscience, 5(7), 700–703.

Schneider, W., Eschman, A., & Zuccolotto, A. (2002). E-Prime: User’s guide. Reference guide. Getting started guide. Psychology Software Tools, Incorporated.

Smout, C. A., Tang, M. F., Garrido, M. I., & Mattingley, J. B. (2019). Attention promotes the neural encoding of prediction errors. PLoS Biology, 17(2), e2006812.

Snodgrass, J. G., & Corwin, J. (1988). Pragmatics of measuring recognition memory: Applications to dementia and amnesia. Journal of Experimental Psychology: General, 117(1), 34.

Team, R. C. (2022). *R: A language and environment for statistical computing*. https://www.R-project.org/.

Vanthornhout, J., Decruy, L., & Francart, T. (2019). Effect of task and attention on neural tracking of speech. Frontiers in Neuroscience, 13, 977.

Wahn, B., & König, P. (2017). Is attentional resource allocation across sensory modalities task-dependent? Advances in Cognitive Psychology.

Xie, Z., Reetzke, R., & Chandrasekaran, B. (2018). Taking attention away from the auditory modality: Context-dependent effects on early sensory encoding of speech. Neuroscience, 384, 64–75.

Yahav, P. H., & Golumbic, E. Z. (2021). Linguistic processing of task-irrelevant speech at a cocktail party. Elife, 10, e65096.

